# Fuzzy set intersection based paired-end short-read alignment

**DOI:** 10.1101/2021.11.23.469039

**Authors:** William J. Bolosky, Arun Subramaniyan, Matei Zaharia, Ravi Pandya, Taylor Sittler, David Patterson

## Abstract

Much genomic data comes in the form of paired-end reads: two reads that represent genetic material with a small gap between. We present a new algorithm for aligning both reads in a pair simultaneously by fuzzily intersecting the sets of candidate alignment locations for each read. This algorithm is often much faster and produces alignments that result in variant calls having roughly the same concordance as the best competing aligners.

---

All Illumina next generation sequencers are capable of producing paired-end reads. These reads represent portions of DNA separated by some distance. Pairing is helpful in aligning the reads against a reference because while organisms often have repetitive genomes, the repetitiveness is less over larger regions, so pairs of reads more often align uniquely than single reads of twice the length [Nakazato]. A typical way to align paired-end reads is first to align one read alone, and then to align the other restricted to locations near to where the first read might align, sometimes repeating the process with the reads reversed and finally choosing the best from the alignment pairs found. BWA-MEM2 [Vasimuddin], Bowtie2 [Langmead], GEM3 [Marco-Sola] and Novoalign [Novocraft] all use some variant of this technique.

We describe the paired-end alignment algorithm implemented in version 2.0 of the SNAP aligner [Zaharia]. Rather than employing a single-end aligner with constraints, it aligns both ends of the pair simultaneously by looking for places where the sets of possible alignments for each end have partial genome matches that are near to one another, which we call *fuzzy set intersection* (“fuzzy” because the locations of the matches do not need to be the same, just to be within the range spanned by a maximally separated read pair). This technique avoids doing expensive evaluations of many candidate alignments that would eventually be dismissed because they are too far from any plausible alignments for the other end of the pair.

SNAP has a hash-table based index of the reference genome which maps fixed-length sequences of bases, called *seeds*, to the set of places they occur in the reference. SNAP chooses several seeds from each read (typically 8 from each) and looks them up in the index. It creates a hit set for each read as the union of the loci where the seeds occur in the reference and fuzzily intersects those sets. It records any loci in the intersection as possible alignment candidates.

Figure 1 is the fuzzy set intersection algorithm and an illustration of its operation, intersecting sets A and B. It starts with the smallest locus *a* from A and binary searches B to find *b*, the smallest locus in B not too small to pair with *a*. If *a* and *b* are close enough to one another to be within fuzzy intersection range, it linearly scans B to find all loci in B that are near enough to *a* and moves *a* to the next element of A; otherwise, it chooses a new *a* by binary searching to find the smallest locus not too small to pair with *b*. It repeats until hitting the end of one of the sets.

**Figure 1:**
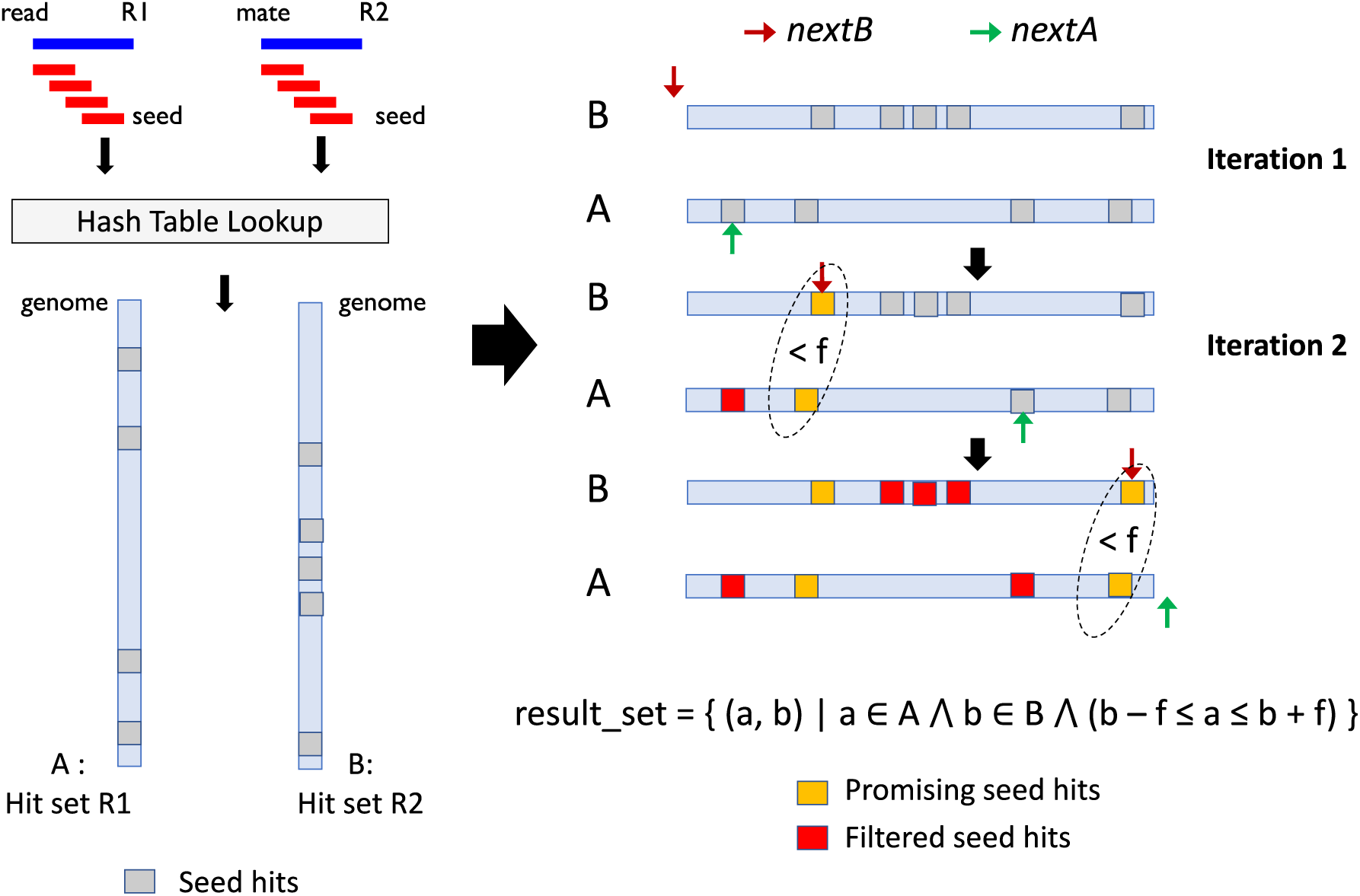
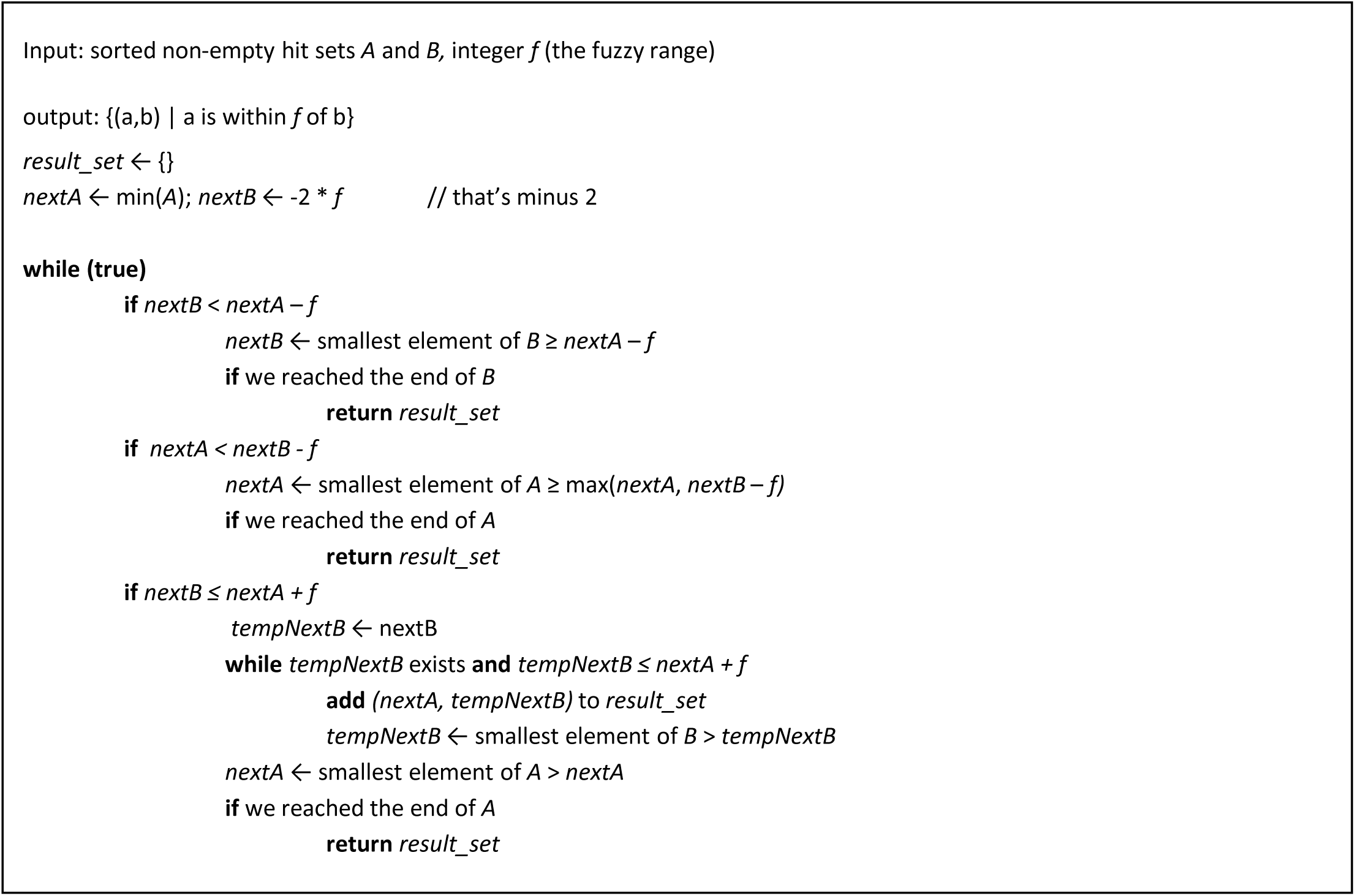
(a) illustrates the fuzzy set intersection algorithm. After the hash-table lookups in seeding, we obtain a sorted set of candidate seed hits for each read: A and B. Starting from the smallest seed hit in set A (indicated by the green arrow, *nextA*), we ping-pong between seed hits in candidate sets A and B using several binary search steps (not shown in the figure). At the beginning of each iteration, the algorithm ensures that all promising candidate seed hits from set A which are less than *nextA* are identified. The algorithm terminates when either *nextA* or *nextB* reaches the end of its respective set. (b) Fuzzy set intersection algorithm pseudocode

SNAP then evaluates the candidates produced by the fuzzy set intersection in much the same way that its single-end aligner does, except that it is evaluating the alignment of both ends of the pair before considering another alignment candidate. The fuzzy set-intersection algorithm could be applied to other aligners, including those that use indices other than a hash table, such as the FM Index [Ferragina] in BWA-MEM [Li 2013], and those that find all possible matches within a given edit distance of the best alignment like GEM.

Eliminating a candidate alignment because it is not in the set intersection is much faster than doing so by scoring it. When the sets are large and the intersection is small, most often the binary search doesn’t need to look at most of the candidates. Conversely, scoring a candidate requires doing some comparison between it and the reference, which involves an algorithm with complexity superlinear in the size of the read such as Ukkonen’s algorithm [Ukkonen] or the banded Smith-Waterman algorithm [Altschul]; in SNAP scoring a candidate is about 10x as expensive as having a candidate in the sets to be intersected.

We measured the size of the hit sets for both reads of every pair in the hg6 sample from the Genome-in-a-Bottle (GIAB) dataset [Zook]. Supplementary Figure 1a shows a heatmap of read pairs by the hit set sizes of each of the reads. While nearly half of the pairs had only one hit for each read it was not uncommon for one read to have few hits while the other had many; 9% of all read pairs had 7 or fewer hits on one read and 128 or more on the other. Supplementary Figure 1b is a heat map of the aligner’s run time divided the same way. The 44% of pairs with only one alignment candidate take 3% of the time, while the bulk of the time is spent on pairs where both reads have many hits.

Figure 2a shows the mean size of the set intersection as a multiple of the **smaller** of the two hit sets for hg6. Because the intersection is fuzzy, this can be larger than one, for instance if the larger set has several hits near to a single hit in the smaller set. Most of these values are substantially less than one, which indicates that the set intersection eliminated many candidate alignments that would need to be evaluated by an aligner even if it perfectly selected the read with fewer candidate alignments to align first. For all the reads together the ratio of the set intersection size to the smaller set size is 0.53, so nearly half of candidate alignments of the read with fewer candidates are eliminated prior to scoring.

**Figure 2:**
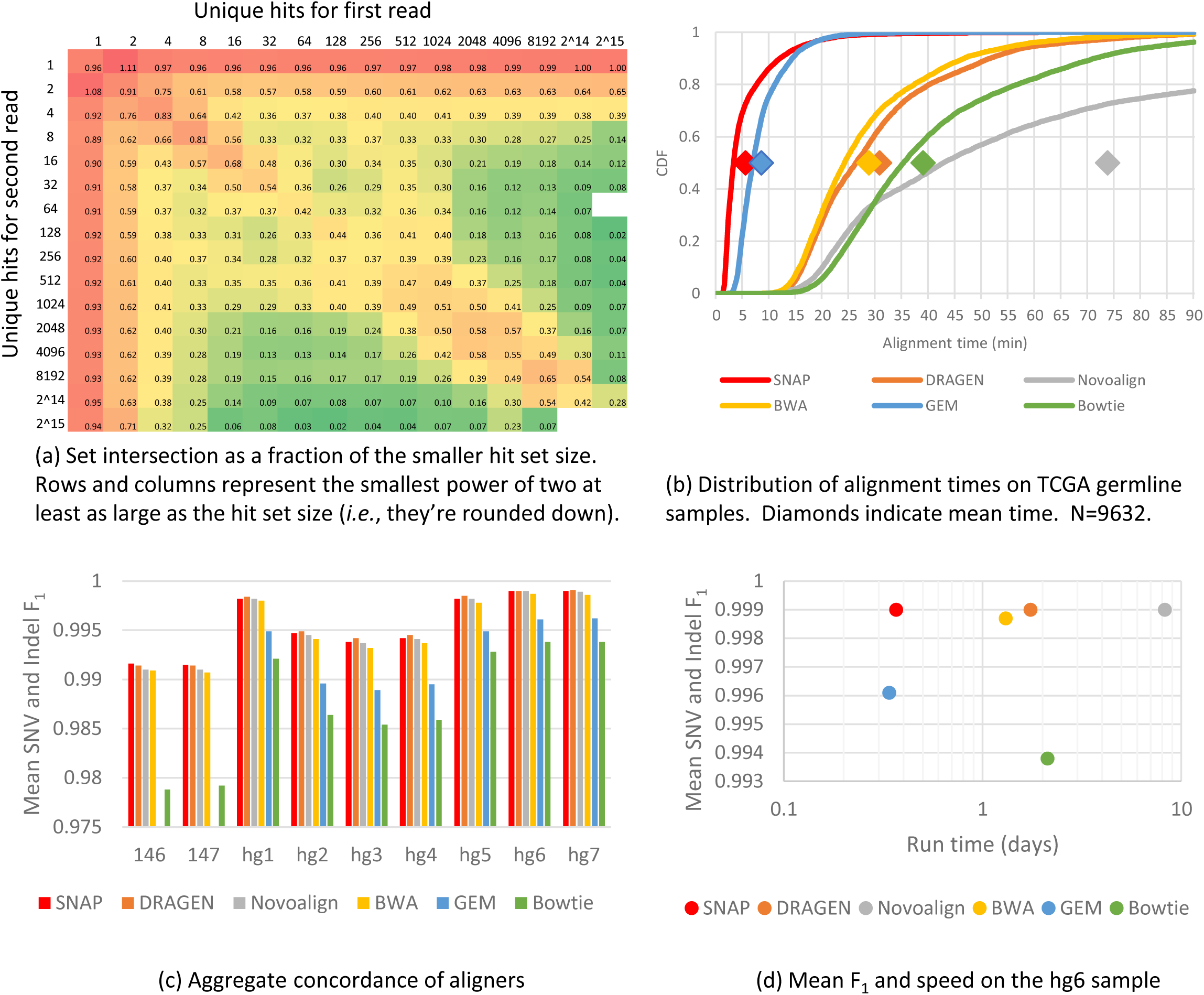
(a) the hg6 size of the fuzzy set intersection divided by the size of the smaller set. It may be greater than one because fuzzy intersection can result in more than one element from the larger set being retained by only one from the smaller. For pairs where the sizes are very different the intersection is commonly substantially smaller than the smaller set, which shows that set intersection eliminates more candidates than just scoring the read with the smaller set first. (b) the CDF of alignment times for five aligners on paired-end germline samples from TCGA. The diamonds show the mean time, which is larger than the median because the distributions have heavy tails. (c) We evaluated the aligners by running a variant calling pipeline using the DRAGEN variant caller and comparing the results against the known truth set provided with the data and plotted the mean F_1_, a composite of the recall and precision for both indels and single nucleotide variants (SNVs). There is no data for 146 & 147 for GEM because it produced output that the downstream tools were unable to use. (d) is a scatter graph showing the mean F_1_ and time to run the aligner (but not the later stages of the pipeline like sorting, duplicate marking and variant calling) for the hg6 sample.

To show its effect on overall run time, we aligned 9,632 paired-end germline samples from The Cancer Genome Atlas (TCGA) using SNAP and five other aligners. SNAP’s mean alignment time for the entire set ranged from 1.5x-13x faster than the other aligners. Figure 2b shows the distribution of run times.

It is difficult to properly evaluate the correctness of an aligner in isolation, because with real reads there is no known ground truth against which to compare, and simulated reads may fail to capture the nuances of real data [Li 2014]. Even if there were ground truth for alignments there would not be for mapping quality (MAPQ) and soft clipping, which can affect the results of analyses that use alignments. Therefore, to determine whether the set intersection algorithm (along with the rest of SNAP) is doing a good job, we evaluated SNAP by using it and other aligners as components in a pipeline that implements one of the most common tasks for sequencing: variant calling human germline samples. We ran each aligner on all seven samples from GIAB (called hg1-hg7) and two from Illumina’s Platinum Genomes [Eberle] dataset (called 146 & 147) and used DRAGEN [Miller] for variant calling. These samples come with high-confidence truth sets. We used hap.py [Krusche] to determine the concordance of the variant calls with the ground truth for single nucleotide variants (SNVs) and insertions/deletions (indels).

The raw results of hap.py are precision and recall for each of indels and SNVs. We use the F_1_ score, the harmonic mean of precision and recall, as a combined metric and then combine the F_1_ scores for indels and SNV as the (arithmetic) *mean F_1_*. SNAP has higher mean F_1_ on all samples than all aligners aside from Novoalign and DRAGEN. Novoalign ties SNAP (to four decimal places) on three with SNAP doing better on the remaining six, while DRAGEN ties on one and is better on six with SNAP having a higher score on the remaining two; SNAP and DRAGEN never differ by more than 0.0004 with mean absolute difference of 0.0002 (Figure 2c). Supplementary Figures 1c-1d provide details on SNV and Indel F_1_ scores. Figure 2d shows both performance and mean F_1_ score for the hg6 sample.

SNAP with the fuzzy set intersection algorithm dominates BWA and Bowtie, having both better performance and better concordance in every test, though BWA’s concordance is close on most. While Novoalign was near to (though very slightly worse than) SNAP in concordance, it had by far the worst performance, 13x slower than SNAP on TCGA and 22x-29x slower on GIAB/Platinum Genomes. GEM was 1.5x slower on TCGA but was overall faster on GIAB/Platinum Genomes with substantially worse concordance on all samples. From this, we conclude that the fuzzy set intersection algorithm improves performance while not compromising (and possibly even helping) concordance.

## Methods

The online methods section provides the details necessary to reproduce our experiments, as well as more details of the results.

SNAP is free software licensed under an Apache 2 license. Its source is available on GitHub at https://github.com/amplab/snap. Its project webpage is https://www.microsoft.com/en-us/research/project/snap/

## Acknowledgements

We would like to thank Novocraft for providing a complimentary license to evaluate Novoalign. Santiago Marco-Sola was very helpful in giving support for running GEM. The results published here are in part based upon data generated by the TCGA Research Network: https://www.cancer.gov/tcga. Richard Karp provided insights that were helpful in SNAP’s design.

## Author Contributions

WB and AS designed the algorithm, wrote the software, designed and conducted experiments and wrote the paper. MZ and RP designed the algorithm and wrote the software. TS and DP designed the algorithm.

## Online methods for “Fuzzy set intersection based paired-end short-read alignment”

This supplemental information for “Fuzzy set intersection based paired-end short-read alignment” is intended to allow the reader to replicate the results in the main paper, as well as presenting more detail and support for these results.

The source and binary code for SNAP is freely available in GitHub at https://github.com/amplab/snap.

Some of the tools used to collate experimental results are available in the “ase” branch of the Git repository.

The spreadsheet with the graphs in the paper and supplementary material together with the scripts that we used to run the experiments are available <u>here</u>^1^. The raw timing and concordance data are available <u>here</u>^2^.

### Computing environment and software versions

We ran all timed experiments on HP Proliant DL360e servers with dual Intel® Xeon® E5-2450L CPUs with hyperthreading disabled. These are 16 core machines. The machines have 192GB of RAM and run Windows Server 2019 Datacenter version 1809. We ran SNAP native under Windows and the rest of the aligners and pipeline tools under Windows Subsystem for Linux (WSL) on Ubuntu 16.04LTS. To get timing results for the DRAGEN aligner we ran its software version (DRAGMAP) on the same machines but did not use the resulting alignments for variant calling.

We used version 3.8 of the DRAGEN variant caller for all variant calls. We ran this on Amazon Web Services (AWS). For the DRAGEN-aligner variant calls, we ran both the aligner (also version 3.8) and variant caller together rather than moving aligned reads from our Windows systems. The performance of the DRAGEN variant caller on AWS is not directly comparable to the others because it runs on different hardware: field programmable gate arrays (FPGAs). The FPGA version of DRAGEN is much faster than the software versions of any aligners we measured, though it is difficult to say exactly how much faster because Illumina’s software combines alignment, sorting, duplicate marking, and variant calling into a single program.

We ran a pre-release version of SNAP version 2.0.0. The differences between the version we ran and the 2.0.0 release should have little or no impact on the results.

The other aligners we used are Bowtie2 version 2.4.1, BWA-MEM2 version 2.0pre2, DRAGEN version 3.8, GEM3 version v3.6.1-25-g82cf-release, and Novoalign V4.02.02 (with Novosort V2.02.00). BWA-MEM chooses between the SSE4.1, AVX2 and AVX512 versions of the source code based on the instruction set supported on the machine. All versions are functionally identical, but the performance of the AVX512 version is ∼1.5x better than SSE4.1. Our machines only support the SSE4.1 instruction set, so we performed all evaluation experiments using SSE4.1.

We used bcftools v1.2 [Li 2009], bedtools [Quinlan] v2.25.0, the Genome Analysis Toolkit (GATK) [Poplin] v4.1.8.0, Hap.py [Krusche] v0.3.12-2-g9d128a9, Java OpenJDK version 1.8.0_252, OpenJDK Runtime Environment build 1.8.0_252-8u252-b09-1∼16.04-b09, and OpenJDK 64-Bit Server VM build 25.252-b09, mixed mode, Picard MarkDuplicates [Picard] version 2.22.4, Python 2.7.12, and samtools [Li 2009] v1.10.

### Input Files

For the concordance experiments we used paired-end data from the Genome-in-a-Bottle (GIAB) project as well as Illumina’s Platinum Genomes. We used all seven samples from GIAB, named hg1 through hg7, and the Platinum Genome samples for NA12877 (ERR194146) and NA12878 (ERR194147) which we refer to as 146 and 147 respectively.

| Sample Name | Read length | Count of Reads | Total Bases |
| --- | --- | --- | --- |
| hg1 | 148 | 6.2G | 918G |
| hg2 | 250 | 883M | 221G |
| hg3 | 250 | 775M | 194G |
| hg4 | 250 | 869M | 217G |
| hg5 | 250 | 4.3G | 1.1T |
| hg6 | 148 | 2.9G | 429G |
| hg7 | 148 | 2.4G | 335G |
| 146 | 101 | 1.64G | 165G |
| 147 | 101 | 1.6G | 162G |

The TCGA samples were all supplied as Binary Alignment/MAP (BAM) files, so we converted them to paired, uncompressed FASTQ^3^ for input to the aligners. First, we filtered out supplementary alignments:

~~~
samtools view -@ 16 -F 2048 -b -o input.filtered.BAM input.BAM
~~~

We then sorted the remaining reads by name:

~~~
samtools sort -n -l 1 -m 10G -@ 16 input.filtered.BAM -T tempBAMFilePrefix -o input.namesorted.BAM
~~~

And used bedtools to produce the FASTQ files:

~~~
bedtools bamtofastq -i input.namesorted.BAM -fq input_1.FASTQ -fq1 input_2.FASTQ
~~~

While SNAP is capable of reading input from a BAM file, we used the same FASTQ inputs as for the other aligners to better facilitate comparison.

The GIAB samples came in multiple compressed FASTQ files. We decompressed them and then concatenated the results to produce a single pair of FASTQ files for each sample.

The GIAB and Platinum samples are all whole genome. Most, but not all, of the TCGA samples are whole exome. The exome is much less repetitive than the rest of the genome, which is at least part of the reason that the TCGA samples aligned much faster per read than the others.

### Reference Genome

We used version 38 of the Genome Reference Consortium human genome reference (GRCh38) from the GATK Resource Bundle (https://storage.cloud.google.com/genomics-public-data/references/hg38/v0/Homo_sapiens_assembly38.fasta). This version is based on the GRCh38 full analysis set (GCA_000001405.15) and includes primary chromosomes, unlocalized scaffolds (_random suffix), unplaced scaffolds (chrUn_ prefix), ALT loci scaffolds (_alt suffix), decoy sequences (_decoy suffix) and HLA sequences (HLA-prefix). The resource bundle also comes with an ALT file specifying the list of ALT contigs for this reference genome (https://storage.cloud.google.com/genomics-public-data/references/hg38/v0/Homo_sapiens_assembly38.fasta.64.alt). Both the reference and ALT contig file are also available as part of Heng Li’s bwakit-0.7.12. In a similar spirit to bwa-postalt.js, SNAP also supports lifting over alignments from ALT contigs to primary contigs as part of its paired-end aligner. For this, an additional ALT-to-primary liftover file (e.g., hs38DH.fa.alt, part of bwakit) is used.

### Index Building

All the aligners use their own index of the reference genome. Other than SNAP and DRAGEN, these indices are FM indices [Ferragina] based on the Burrows-Wheeler Transform (BWT) [Burrows], which produces a smaller but somewhat less performant index than SNAP’s hash table. Building an index is something that needs to be done only when the reference genome changes, not on every alignment. In the following command lines “*index*” (in italics) is the name of the index generated by the aligner’s indexing function. BWA imputes the index from the name of the reference FASTA.

We built the indices on one machine and then copied them to the machines where we made the measurements. This copy reduced or eliminated the disk fragmentation that may have existed from the initial index builds, which probably reduced the index loading time during the measurement runs. We also often did multiple consecutive runs, which may cause the indices to be in the operating system cache which would also result in faster index loading time. The index loading times typically were on the order of seconds to a few minutes (when the indices weren’t in the OS cache), which is small relative to the GIAB and Platinum alignment times. The TCGA runs were done one after another, so all but the first on a given server would likely have had the index in the OS cache.

We used the following commands to build the indices:

~~~
snap.exe index Homo_sapiens_assembly38.fasta *index* -large -altLiftoverFile
Homo_sapiens_assembly38.fasta.alt
~~~

We used a “large” SNAP index, which trades memory for speed. For this particular reference genome it increased the size by about 30%. Since the machines we used all had sufficient memory to store the larger index, this seemed worthwhile.

For BWA, the command to create the index is simple:

~~~
bwa-mem2 index Homo_sapiens_assembly38.fasta
~~~

For Bowtie, we used:

~~~
bowtie2-build Homo_sapiens_assembly38.fasta *index* --threads 16
~~~

For GEM, we used:

~~~
gem-indexer -i Homo_sapiens_assembly38.fasta -o *index* -t 16
~~~

For Novoalign we used:

~~~
novoindex *index* Homo_sapiens_assembly38.fasta
~~~

For DRAGEN we used:

~~~
dragen-os --build-hash-table true --ht-reference hg38.fa --output-directory
*index* --ht-num-threads 16 --ht-alt-liftover bwa-kit_hs38DH_liftover.sam
~~~

### Aligning

We used the following command lines for the aligners. For SNAP, we used:

~~~
snap.exe paired *index* input_1.FASTQ input_2.FASTQ -o output.BAM -so -sm 20 -
hc- -eh -d 20
~~~

SNAP by default adds read group information to the output Sequence Alignment/MAP (SAM) records if not provided by the user on the command line. Also, we used the default values of minimum template length (0) and maximum template length (1000) for paired-end reads for all of the evaluated datasets.

For Bowtie we used:

~~~
bowtie2 -t -x *index* -t –maxins 1000 -p 16 -1 input_1.FASTQ -2 input_2.FASTQ -
S output.SAM -t --rg-id 4 --rg LB:X --rg PL:ILLUMINA --rg SM:20 --rg PU:unit1
~~~

For BWA we used:

~~~
bwa-mem2 mem -Y -K 100000000 -R
’@RG\tID:4\tLB:$1\tPL:ILLUMINA\tSM:20\tPU:unit1’ -t 16 reference.FASTA
input_1.FASTQ input_2.FASTQ -o output.SAM
~~~

For GEM we used:

~~~
gem-mapper -I *index* -1 input_1.FASTQ -2 input_2.FASTQ -o output.SAM -M 1 -t
16 -r ’@RG\tID:4\tLB:$1\tPL:ILLUMINA\tSM:20\tPU:unit1’
~~~

GEM has fast and sensitive modes. We ran it in fast mode because sensitive mode crashed on every input we tried. For the Platinum Genome samples, GEM produced output that both the standard pipeline and Novosort were unable to process, so we did not obtain those results.

For Novoalign we used:

~~~
novoalign -d index -f input_1.FASTQ input_2.FASTQ --alt on -o BAM 0
"@RG\tID:4\tLB:$1\tPL:ILLUMINA\tSM:20\tPU:unit1" > output.unsorted.BAM
~~~

Our machines did not have enough memory for Novoalign to align the hg1 and hg5 samples. Therefore, we split each of them into ten equally sized pieces, aligned those pieces, combined the resulting SAM files and used the consolidated SAM files as input to the standard (not Novosort) pipeline described below. We did not include dividing and combining the input and output files in Novoalign’s execution time.

For DRAGEN we used:

~~~
dragen-os -r /mnt/d/sequence/indices/dragen-38 -1 input_1.FASTQ -2
input_2.FASTQ –output-directory /mnt/d/temp –output-file-prefix
input.dragen3.8-software
~~~

We measured the run time of all aligners other than SNAP by noting the time immediately before and after the aligner process run. SNAP combines alignment with sorting, duplicate marking and indexing in the same process so we used the sum of its reported index load and alignment times.

### Post-processing of aligner output

To turn the output of Bowtie, BWA and GEM into sorted, indexed, duplicated-marked BAM files we ran a pipeline of standard tools. These aligners all produce SAM files. We first converted the SAM files into unsorted BAM files using samtools. Recall that the machines running this pipeline have 16 cores.

~~~
samtools view -@ 16 -S -1 -b output.SAM > output.unsorted.BAM
~~~

Next, we sorted by aligned coordinate:

~~~
samtools sort -@ 16 -m 10G -l 1 output.unsorted.BAM -o output.sorted.BAM -T tempDirectory
~~~

Then, we used Picard MarkDuplicates to mark duplicates and create the final BAM file:

~~~
java -jar picard.jar MarkDuplicates I=output.sorted.BAM O=output.BAM
M=/dev/null 2>&1 >> /dev/null
~~~

Finally, we indexed the BAM file:

~~~
samtools index -@ 16 output.BAM output.BAM.BAI
~~~

Novoalign produces (unsorted, unmarked) BAM files as its output and has its own tool, Novosort, for converting them into sorted, duplicate marked BAMs.

~~~
novosort –md output.unsorted.BAM > output.BAM
~~~

We indexed the resulting BAM with the same samtools command we used for other (non-SNAP) aligners:

~~~
samtools index -@ 16 output.BAM output.BAM.BAI
~~~

SNAP produces sorted, duplicate-marked, indexed BAM files directly, so we didn’t run any additional tools with SNAP.

For variant calling we used the FPGA version of the DRAGEN aligner on AWS that is integrated with the DRAGEN variant caller. For measuring alignment time, we used the software version of DRAGEN aligner (DRAGMAP), but did not run later pipeline steps because a) the software version produces the same results as the FPGA version and b) DRAGEN will presumably eventually come with its own tools, so measuring the performance of standard pipeline after alignment is not useful. Therefore, DRAGEN is omitted from the whole pipeline performance comparisons.

### Variant calling

We used version 3.8 of the DRAGEN variant caller running on AWS for variant calling. For aligners other than DRAGEN, we first aligned, sorted, and marked duplicate reads as described above and then loaded the resulting BAM and BAM Index (BAI) files into AWS’s S3 storage service. We had to provide a large value for DRAGEN’s --bin-memory parameter because otherwise it failed on some of the inputs.

We then ran the DRAGEN variant caller as follows:

~~~
dragen -f -r ∼/index -b s3://dragentest/*input*.bam --enable-variant-caller
true --output-directory /data/dragen-output/ --output-file-prefix *input* --
enable-map-align false --enable-sort false --bin_memory 90000000000
~~~

For the DRAGEN aligner, we loaded the input FASTQ files into S3 and ran:

~~~
dragen -f -r ∼/index -1 s3://dragentest/*input*_r1.fastq.gz -2
s3://dragentest/*input*_r2.fastq.gz --enable-variant-caller true --output-
directory /data/dragen-output/ --output-file-prefix *input*.dragen3.8 --enable-
map-align true --enable-sort true --RGID foo --RGSM bar --RGPL foo --RGPU bar
--enable-duplicate-marking true --enable-map-align-output true --bin_memory
90000000000
~~~

### Measuring concordance

We used hap.py to evaluate concordance between the variant calls from the various aligners and the known truth sets for each sample. First, we did some preprocessing steps on each Variant Call Format (VCF) file:

~~~
bcftools annotate -x FORMAT/AD -O z -o *output*.NoAd.VCF.gz *output*.VCF
bcftools view -c 1 -O z -o *output*.NoAd.NoHomRef.VCF.gz *output*.NoAd.VCF.gz
~~~

Then we ran hap.py and saved the output:

~~~
python hap.py ground-truth.VCF.gz *output*.NoAd.NoHomRef.VCF.gz -r
reference.FASTA -o output.concordance --engine=vcfeval -f ground-truth.BED
tar cvf *output*.concordance.TAR .
~~~

### Heat maps

We generated the heat maps in Figure 2 and Supplementary Figure 1 by adding instrumentation to SNAP’s code. That instrumentation is available in the SNAP sources. To enable it, change the value of INSTRUMENTATION_FOR_PAPER in SNAPLib/AlignerOptions.h to 1 and rebuild SNAP. Running the version with instrumentation will produce a file called SNAPInstrumentation.txt in the directory in which you run the program.

The same file contained the data to support the claim in the main paper that the hg6 aggregate set intersection was 0.53 of the smaller sets. This was done by counting the total size of the set intersections across all the read pairs in hg6 and dividing that by the sum of the sizes of the smaller sets. These two numbers are in the instrumentation file.

The colors in the heat maps were generated by Microsoft Excel.

### Timing error bars

Other than Novoalign and Bowtie, we ran each of the GIAB and Platinum Genome alignment tests (but not the rest of the pipeline) five times and computed the standard error. We did not compute error bars for the full pipeline. All the error bars were less than a half percent of the overall run time aside from DRAGEN on 146 (1.2%) and BWA on hg2 (1.5%).

### Detail on Results

The main body of the paper presents a summary of the results. Here we present more detail. For a given concordance run, every variant is either called or missed, and every variant call is either correct or incorrect. This gives rise to counts of true positives (TP) (variants that were correctly called), false positives (FP) (variant calls that do not correspond to variants) and false negatives (FN) (variants that were not called). One could also imagine true negatives, which are all the variants that could have been called but were not, but we do not use them in our analysis; they would greatly outnumber the other categories.

Precision is TP/(TP+FP) and roughly measures lack of false positives. Recall is TP/(TP+FN) and roughly measures how likely a variant was to be called. The F_1_ score is the harmonic mean of precision and recall, F_1_ = 2 * (precision + recall) / (precision x recall). Finally, we computed a composite score, the mean F_1_, by taking the arithmetic mean of the SNV and indel F_1_ scores: mean F_1_ = (F_1recall_ + F_1precision_) / 2.

Supplementary Figure 2 shows the SNV and Indel precision and recall for each of the aligners on each of the GIAB and Platinum Genome samples.

Overall, the precision and recall of the samples for most aligners is quite good. Supplementary Figure 3 makes clear small differences of values close to 1 by showing the mean F_1_, SNV and Indel “9s”, which is -log_10_(1-score) for the relevant score. Informally, it is counting the number of 9s at the beginning of the decimal value, so for example 0.9 is 1 and 0.99 is 2.

Supplementary Figures 4 and 5 are scatter graphs like Figure 2d, but for the 8 samples not shown in the main paper. When reading these graphs, it’s worth keeping in mind the scales. The range of mean F_1_ is usually less than 1% from worst to best and always less than 1% when ignoring Bowtie (though a 1% difference can be quite meaningful particularly for values as close to 1 as these), while the time axis is on a log scale covering three orders of magnitude. The nearest the fastest and slowest aligners get to one another is 24-fold, and the farthest is 39-fold.

### Reverse Complements

DNA is double stranded with each strand being the reverse complement of the other: it goes backwards and has complementary bases (T pairs with A and C with G). In paired-end reads one end is from the forward strand and the other end is from the reverse complement, though it’s not possible to tell which is which *a priori*. SNAP runs the entire set intersection twice for each read pair, once assuming the pair is FR (Forward-Reverse) and once assuming RF. We skipped this detail in the main paper and elsewhere in the supplementary material for clarity.

SNAP’s index only includes the forward version of the reference, so SNAP alters the read that it assumes is the reverse complement before looking it up to obtain the set to use in the intersection operation. For example, in the FR case, SNAP would align the first read and the reverse complement of the second.

It is possible that a read pair is FF or RR if the pair spans a structural variant such as inversion. In this case, SNAP’s paired-end aligner will fail to find an alignment and it will fall back to aligning each of the reads in the pair individually with the single-end aligner.

### Other properties of SNAP

While not the main point of this paper, SNAP has several optimizations other than the set intersecting candidate selection algorithm. Understanding them makes it easier to grasp SNAP’s overall behavior. These optimizations are present in both SNAP’s paired-end and single-end aligners.

The first optimization is using Landau and Vishkin’s algorithm [Landau] for scoring. This algorithm computes an edit distance (the number of mismatched bases or single base insertions or deletions) between the read and reference if the total distance is less than some value *k*. Because the cost of the computation depends on the smaller of the edit distance between the two and *k*, highly similar read and reference sequences can be matched quickly and keeping *k* small can efficiently discard bad alignment candidates. The insight here is that there is no need to know the true edit distance for a candidate alignment if it is much larger than the best candidate that has already been found (or, if finding the best *n* candidates, the *n*th best that’s been found). Candidates slightly worse are useful for computing the mapping quality, but beyond that finding the exact edit distance is a waste of time since the result will be discarded.

Since the maximum edit distance that will be computed depends on the best candidate alignment found so far, it would be optimal to score the overall best candidate alignment first, thus reducing *k* and improving performance. Of course, doing this requires knowing which alignment candidate is best, which would obviate the need for the rest of the alignment process and so is unrealistic. Instead, SNAP uses a heuristic to try to score better candidates earlier. It is to count the number of seed hits that support a particular alignment candidate and to score the candidates in the order of count of hits. That is, if SNAP looked up *x* seeds it will first score the candidate alignments supported by all *x* of them, then by *x-1*, and so on. We instrumented SNAP running hg6 and found that 14% of pairs had more than one candidate alignment scored. 46% of those scored the best candidate first.

As others have observed [Ahmadi], the number of misses in any disjoint seed set is also a lower bound on the eventual edit distance because each miss must be caused by at least one difference between the read and the reference [Baeza-Yates]. Because the seeds are disjoint, the differences must all be different from one another. If that lower bound is high enough that the read could not possibly affect the result of the alignment (*i.e*., it’s larger than *k*), then SNAP skips scoring it altogether. This optimization is why the number of read pairs that have only one candidate location scored is higher than the fraction of pairs that have only one hit on each read; after scoring the first candidate, the other(s) sometimes are eliminated without having to score them. Supplementary Figure 1a shows 44% of the pairs have only one candidate scoring location, while the instrumentation described in the preceding paragraph showed that 86% of reads have only one candidate location scored.

Using simple edit distance to choose the best alignment does not work well for alignments that have insertions or deletions of more than one base. Edit distance will sometimes prefer candidate alignments with more indels but having fewer total bases as part of indels, which is biologically less likely. Consider for example aligning read GAAAATATAAAA against reference GAAAAAAAAA. The alignment with two single-base insertions of T has an edit distance of 2, while the alignment with a three base insertion of TAT has an edit distance of 3, and so the edit distance metric will prefer the two small insertions while the three base insertion is more biologically plausible.

Affine gap scoring allows the cost of indels to vary disproportionally to their length. However, it is much more expensive to compute than edit distance. SNAP handles this by using edit distance to eliminate candidate alignments where it can prove affine gap would not produce a different result. After scoring all the reads using edit distance, it takes the remaining alignment candidates where affine gap might make a difference and computes their affine gap score to produce the final result. SNAP vectorizes affine gap scoring using Farrar’s striped algorithm [Farrar]. Farrar’s algorithm stripes adjacent query characters across different SIMD vectors to simplify data dependencies in the dynamic programming formulation. Farrar’s algorithm computes the entire dynamic programming matrix and has complexity O(MN / W), where M is the reference text length, N is the read pattern length and W is the width of SIMD vector (e.g., 128-bit SSE can compute 8 cells in parallel, assuming 16-bit precision for scoring). However, computing the entire matrix is not necessary for most of the candidate reference locations, since typically after scoring the first few locations, based on Landau & Vishkin’s algorithm, SNAP reduces *k*, the maximum edit distance limit for scoring subsequent locations. To reduce computation with smaller maximum edit distance limits, we implement a banded version of Farrar’s fully vectorized implementation in SNAP. The banded version only computes cells that lie within a 2k + 1 band of the major diagonal of the dynamic programming matrix, where k is the current maximum edit distance limit for any alignment. SNAP invokes the banded affine gap implementation only when the length of the query pattern to be aligned against the reference is of length >= 3 * (2k + 1). At smaller query lengths, the fully vectorized implementation provides better performance. This is because when the query length is small, there are fewer mis-speculations in the Farrar algorithm for the fully vectorized implementation.

When SNAP is unable to find an alignment with the paired-end aligner, or when the alignment it finds is of sufficiently poor quality, it aligns each of the reads from the pair separately with its single-end aligner. This is necessary for paired-end reads that span a structural variation, for example. It also happens in the case of reads that do not align at all. For the GIAB and Platinum samples the rate of unaligned reads (excluding those that were too short or contained too many Ns) ranges from 0.4% for hg4 to 1.6% for hg5. The fraction of reads eventually successfully aligned by the single-end aligner is also low, ranging from 1.4% for hg4 to 3.1% for 147.

Version 38 of the Genome Reference Consortium’s human reference contains alternate (ALT) contigs. These are regions of the genome where there are significant differences within the population. One haplotype is chosen to be the primary for a chromosome and any others are assigned to ALT contigs. These ALT contigs are often very similar to the primary. If an aligner did not have special support for them, it might decide that a read would map well to both the primary and to an ALT contig and so give low mapping quality to whichever alignment(s) it chose to report. This low mapping quality might then cause downstream tools to incorrectly ignore (or give low weight to) the read and therefore lose its information. SNAP understands which contigs are ALTs, prefers alignments to the primary, and does not consider ALT candidate alignments when computing mapping quality for alignments to the primary. DRAGEN, BWA and Novoalign have similar support, while GEM and Bowtie do not. This may in part explain GEM and Bowtie’s lower concordance than the ALT-aware aligners.

When SNAP finds a mapping to an ALT contig that is much better than any mapping to a primary contig it will attempt to lift over the mapping from one to the other. That is, it will use information provided in the ALT liftover file that came with the reference to determine what locus in the primary contig most closely corresponds to the portion of the ALT to which the mapping was found, and then map the read there.

In addition to aligning reads, SNAP implements the rest of the standard pipeline to create a finished BAM file. It sorts aligned reads by their aligned coordinates, marks duplicates, compresses them into the BAM format and indexes the resulting BAM file. Doing it this way rather than having separate tools for each step is more efficient because it avoids reading and writing the typically very large data files multiple times. There is at most one additional trip to disk if memory is too small to hold the entire data file when SNAP is producing sorted output. SNAP aligns reads into batches (with size determined by available memory), sorts those batches in memory and then writes them to disk. Once all reads are aligned it re-reads the batches and uses the standard external single-stage merge-sort algorithm [Knuth] to create a final sort. While this is being produced it looks for duplicate reads to mark, and then compresses and writes the final output.

Supplementary Figure 6a shows the time for each of the GIAB/Platinum Genome samples to go from the input FASTQ to the final sorted, duplicate marked, indexed BAM for aligners other than DRAGEN. Bowtie, BWA and GEM produce SAM output and use the standard pipeline for post-processing. The MarkDuplicates code in the Picard tools is particularly slow and contributes the bulk of the time other than alignment. Novoalign uses its custom Novosort tool, which is much faster than the standard pipeline and in some instances is also faster than SNAP. Supplementary Figure 6b shows the distribution of full pipeline time for the TCGA samples; the superior performance of Novosort causes the Novoalign pipeline to be faster than BWA and Bowtie on average, even though the Novoalign aligner is 2.6x and 1.9x slower than BWA and Bowtie respectively.

### SNAP parameter settings

We chose SNAP’s default seed size by trying different values on the hg3, hg7 and 147 samples. We used these samples because they spanned the range of read lengths in our dataset: 250, 148 and 101 respectively. Supplementary Figure 7 shows the results. Each graph shows points with speed and concordance for each seed size we tried. The lines connect the points in order of seed size. In all cases the smallest size we tried was 19 and it resulted in the slowest (rightmost) performance. The general trend is clear: larger seed sizes improve performance and have an optimum for concordance at an intermediate value. That said, the difference in concordance based on seed size is quite small, while the performance effects are not, so it might make sense for many applications to use larger than default seed sizes. We chose 24 as the default seed size for SNAP.

Supplementary Figure 7d shows the range of variation for varying SNAP’s seed size on hg7 put in the context of the other aligners. Unlike Supplementary Figures 7a-c the time axis is on a log scale. It graphically illustrates the point that the performance effects are large while the concordance changes are relatively small.

SNAP has numerous other parameters whose default values remain unchanged from SNAP v1.0. When choosing the default values for 1.0 we did a similar process to that for seed size, except that we used Haplotype Caller [Poplin] for variant calling. However, unlike seed size we did not see large changes in either performance or concordance for reasonable values of other parameters. Therefore, we did not repeat the experiments for v2.0.

### Uses of SNAP

Versions of SNAP have been available since 2011, and it is widely used in the community. It has been used in dozens of research studies, often in the context of metagenomics, where it may be used to filter out reads from contaminants such as human or other host DNA [Dash; Greininger; Guo; Henriques; Joyjinda; Li 2018; Masembe; Sorek] or in any of a number of ways to identify microbes or validate results [Birdsell; Bouquet 2017; Bouquet 2019; Brown; Dias; Fortney; Franzke; Knight; Lees; Onimaru; Rahman; Stroehlein; Wesolowska], sometimes as a component of SURPI or TransRate [Smith-Unna]. Other studies use it to align reads against a human or other reference [Dodman; Pearl; Readhead; Woronik], or as part of building a reference for a new species [Huang 2020; Low; Mamrot; Teh; Thorpe]. It is also a component of several bioinformatics frameworks and tools [Banerjee; Byma; Folarin; Gou; Huang 2018; Lin; Magis; Sahl; Tithi]

---

**Supplementary Figure 1:**
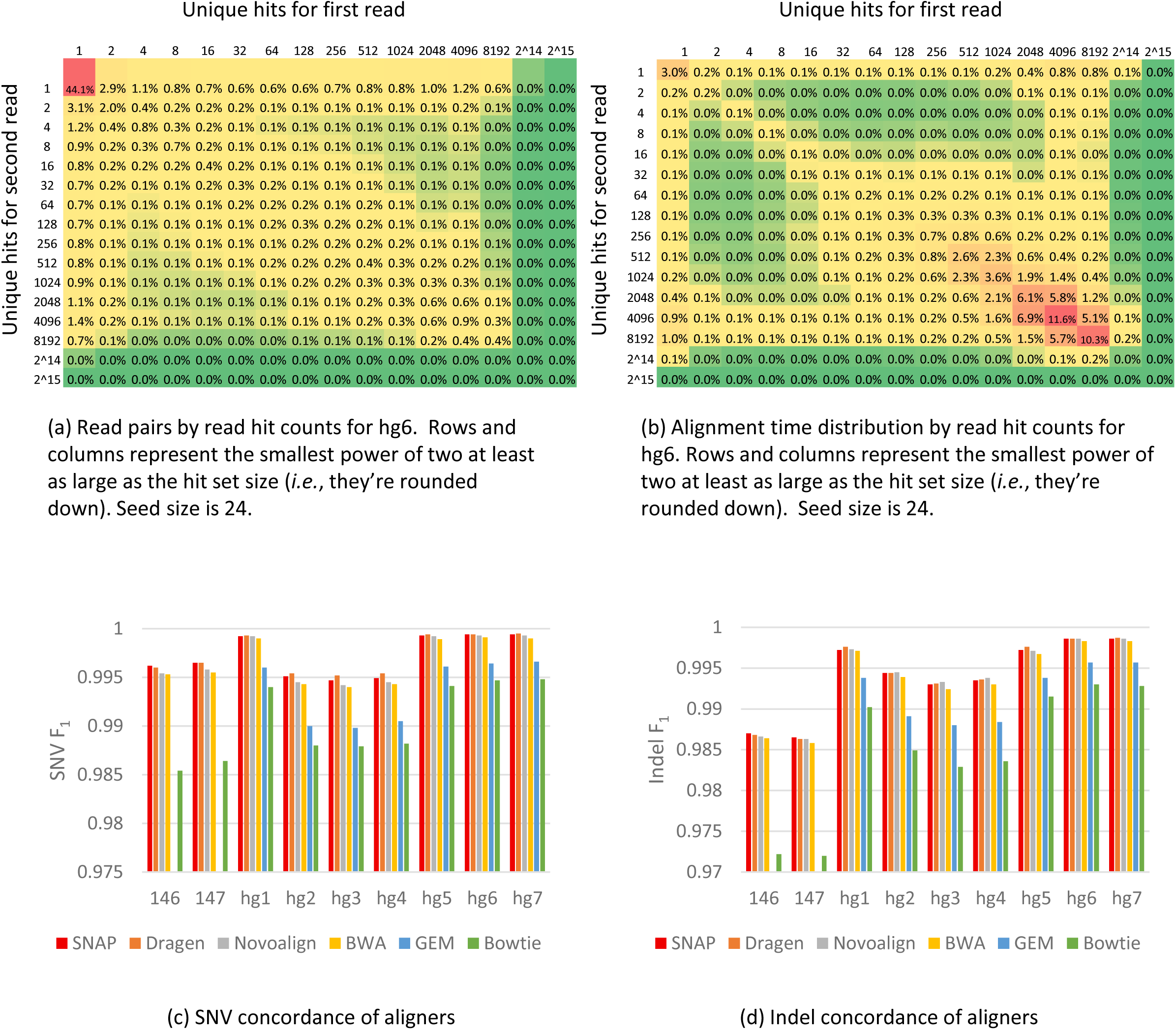
(a) the distribution of read pairs in the GIAB hg6 sample divided by the number of unique genome locations found by the seeding algorithm for each of the reads. Slightly more than half have 1 or 2 candidate locations, but a substantial number have many. It is common to have many hits on one seed but few on the other, which shows that the set intersection often will be much smaller than the larger set. (b) shows the hg6 distribution of time spent by the hit counts for the two reads, showing that well over half is consumed by pairs with many hits on both seeds (> ¾ of the time for pairs with at least 512 hits on each read), but very little by pairs with many hits on exactly one seed. (c) and (d) display the SNV and Indel F_1_ scores respectively for all sample-aligner pairs. (c) and (d) do not have results for GEM for 146 and 147 because it produced output that the downstream tools were unable to use.

**Supplementary Figure 2:**
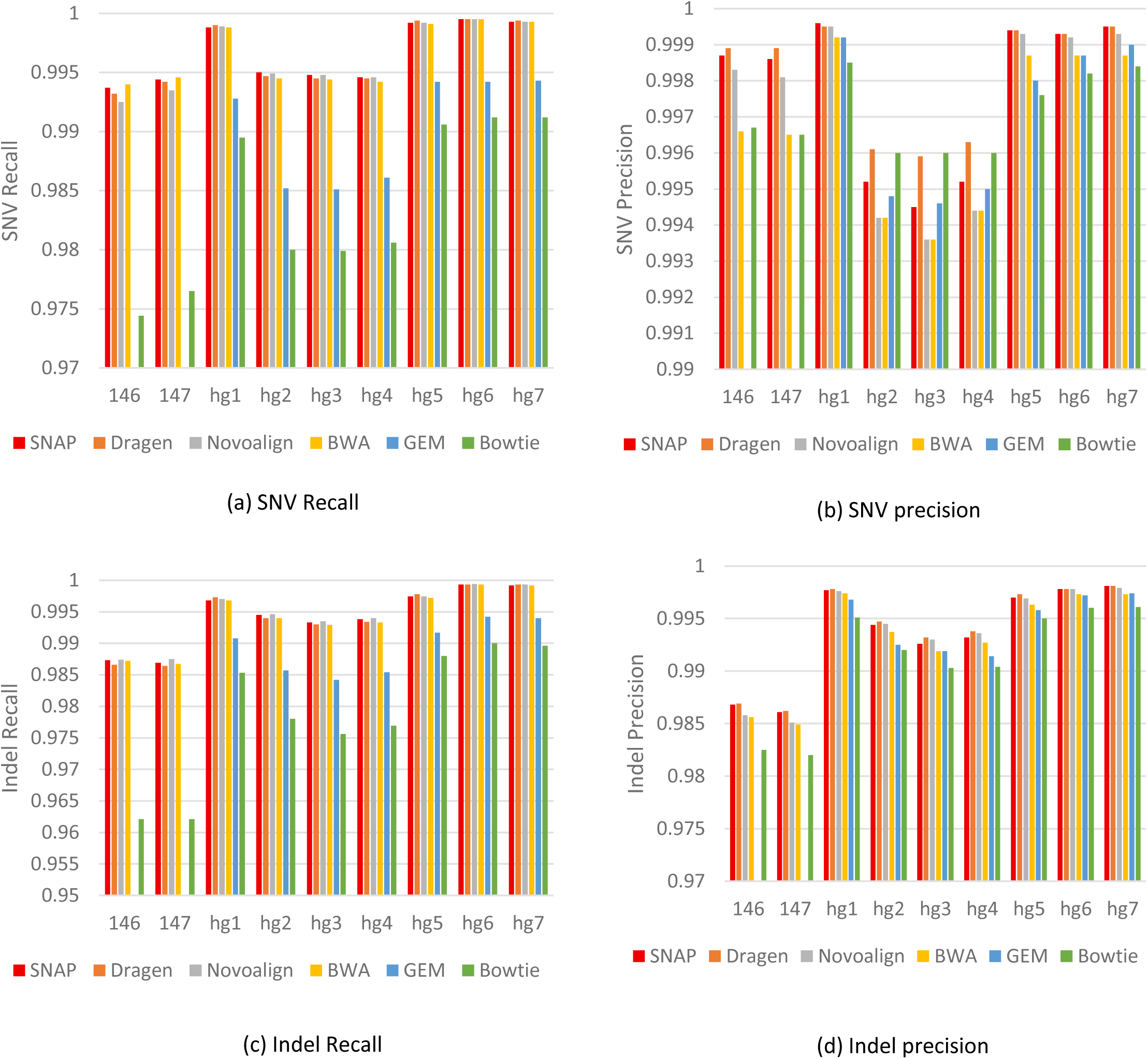
The four components of the mean F_1_ score for each of the aligners on each of the samples for which we have ground truth. GEM did not produce results for 146 and 147 that the downstream tools could use.

**Supplementary Figure 3:**
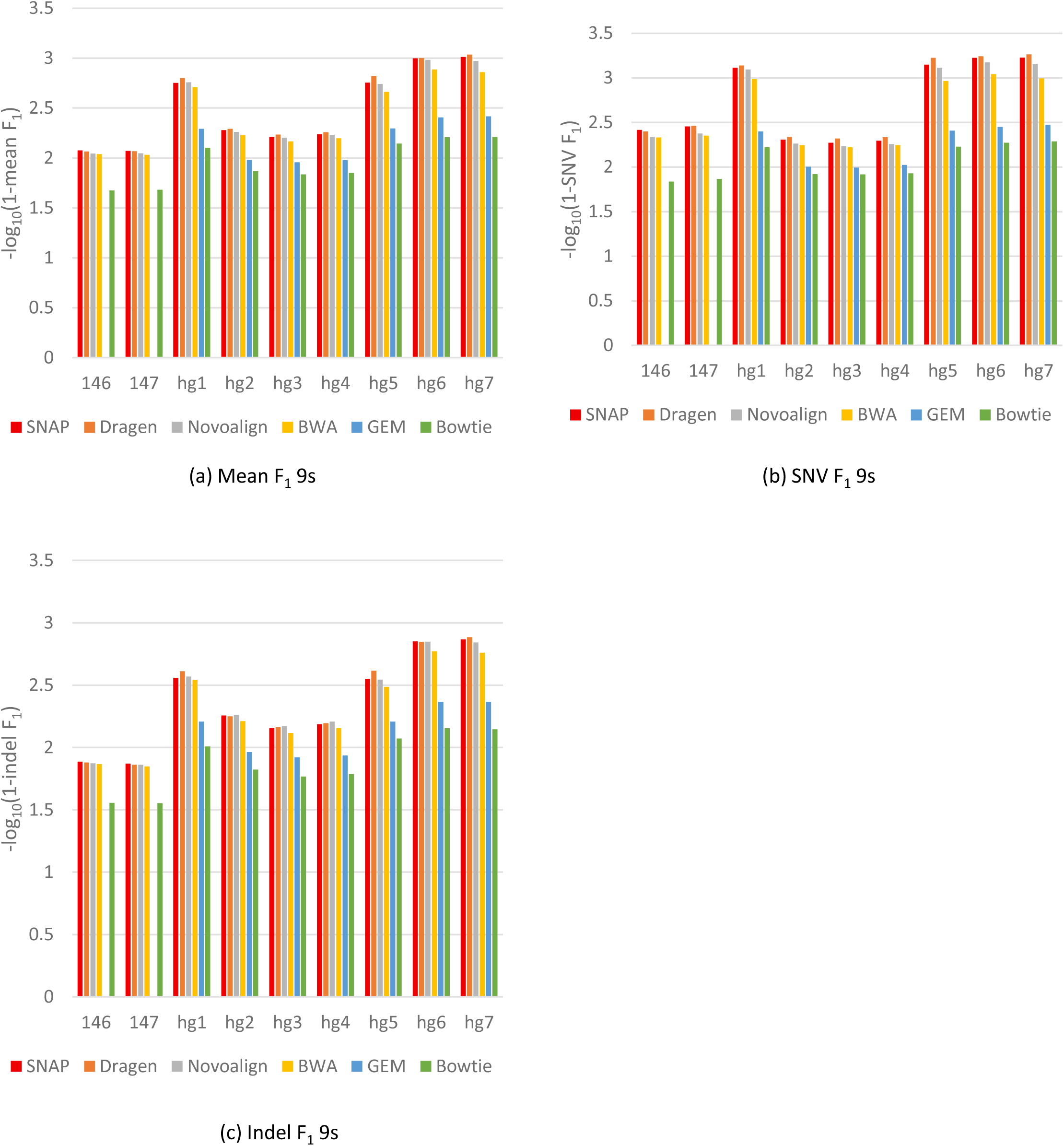
Concordance results expressed in 9s. GEM did not produce results for 146 and 147 that the downstream tools could use.

**Supplementary Figure 4:**
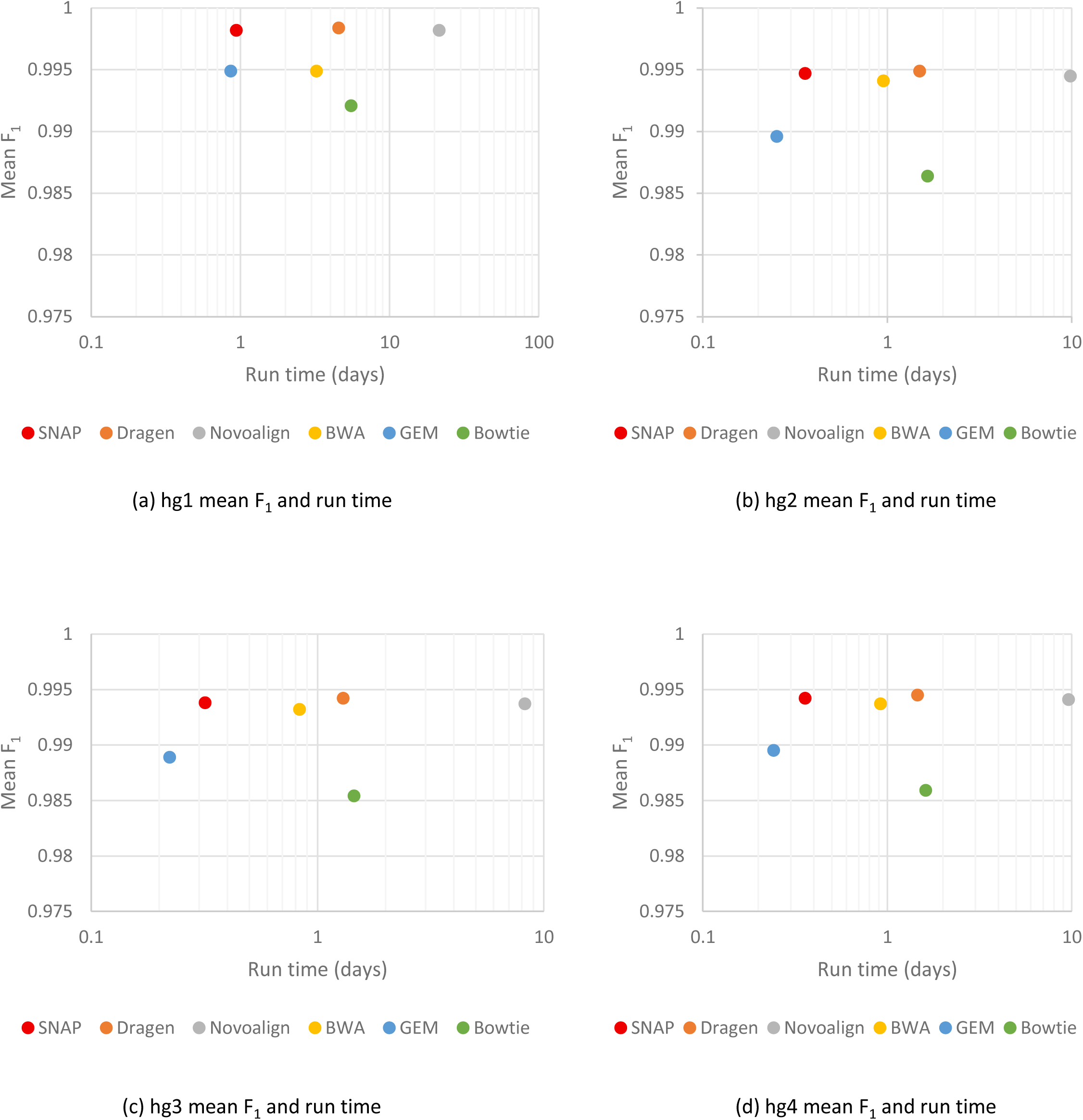
mean F_1_ concordance and aligner run time for some GIAB samples.

**Supplementary Figure 5:**
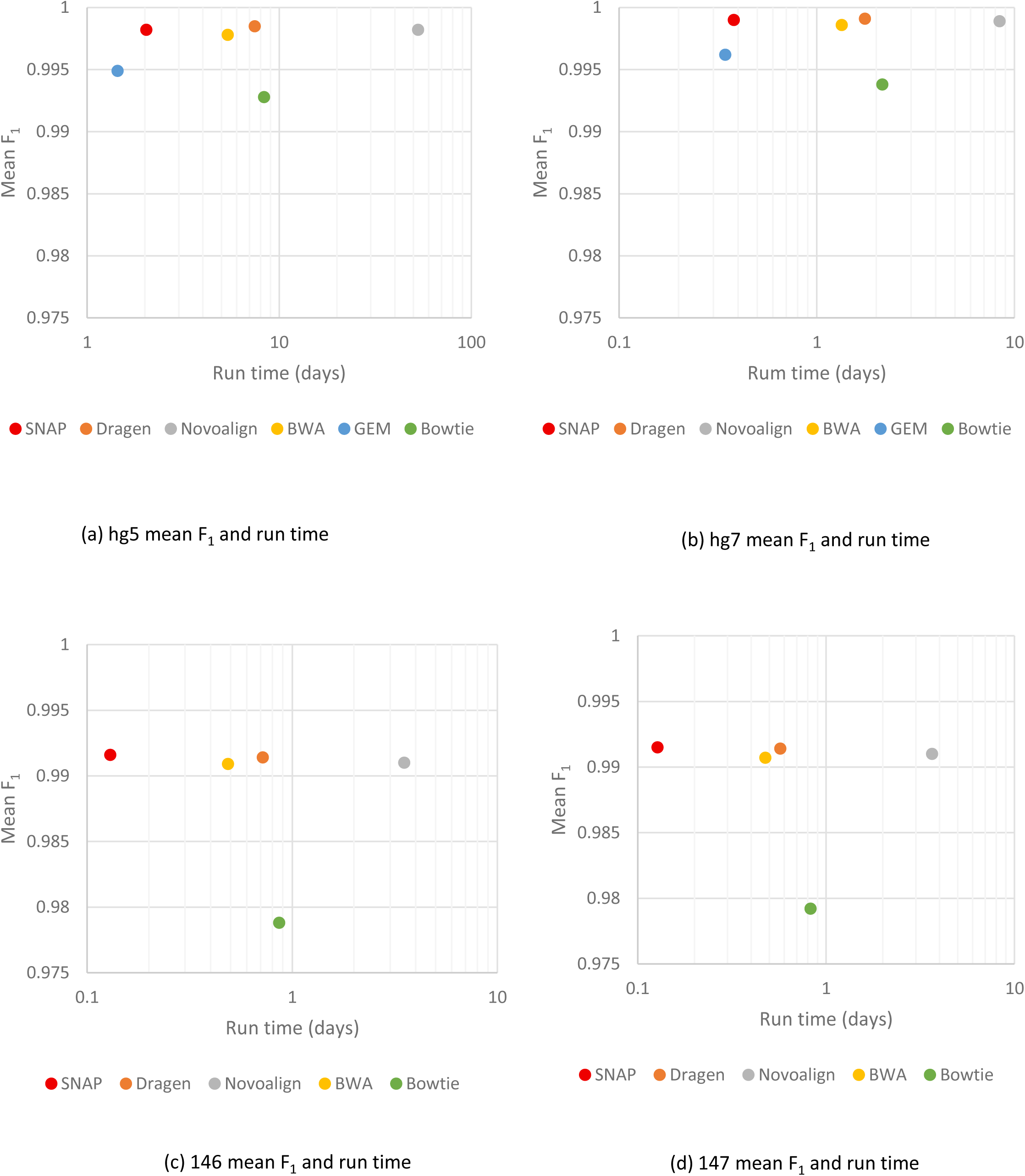
mean F_1_ concordance and aligner run time for some GIAB and Platinum Genomes samples. The Platinum Genomes samples do not have results for GEM because the rest of the pipeline could not process its output. Hg6 is omitted because it is included in the main paper as figure 2b.

**Supplementary Figure 6:**
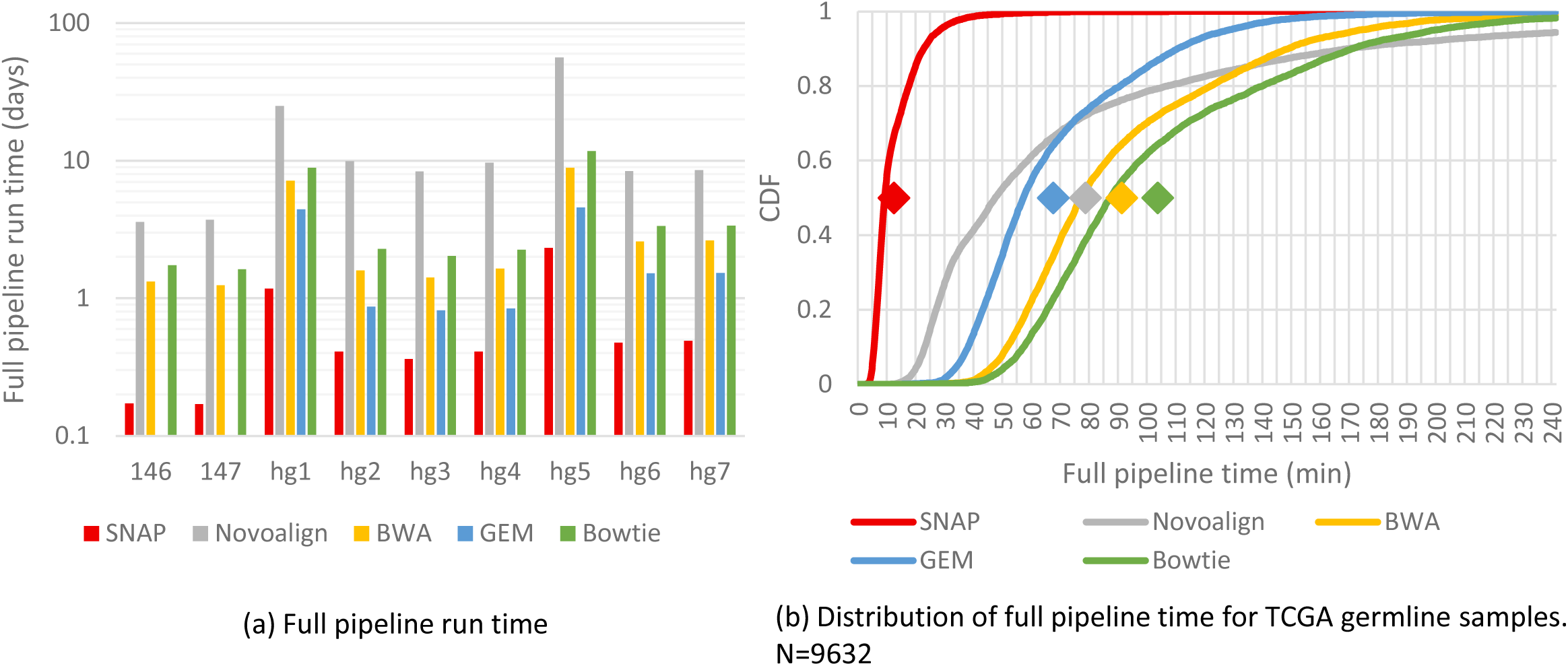
Run times of the full pipeline from FASTQ to sorted, indexed, duplicate-marked BAM file. (a) GIAB and Platinum Genome Samples. (b) TCGA germline samples. SNAP uses its internal pipeline for all samples. BWA, GEM and Bowtie use a standard samtools and Picard pipeline; the bulk of the time was consumed by Picard’s MarkDuplicates code except for BWA and Bowtie on hg6, which used Novosort since the standard pipeline crashed. Novoalign used the Novosort tool on all samples other than hg1 and hg5, on which it used the samtools/Picard pipeline. It is notable that in the smaller, often whole exome TCGA samples that the difference in performance between Novosort and samtools/Picard is more than large enough to make up for Novoalign’s much slower alignment performance, though it does not in the GIAB and Platinum Genome samples. DRAGEN is not included because a software (*i.e.*, not FPGA) version of its pipeline is not available. GEM did not produce output that could be consumed by the rest of the pipeline for 146 and 147.

**Supplementary Figure 7:**
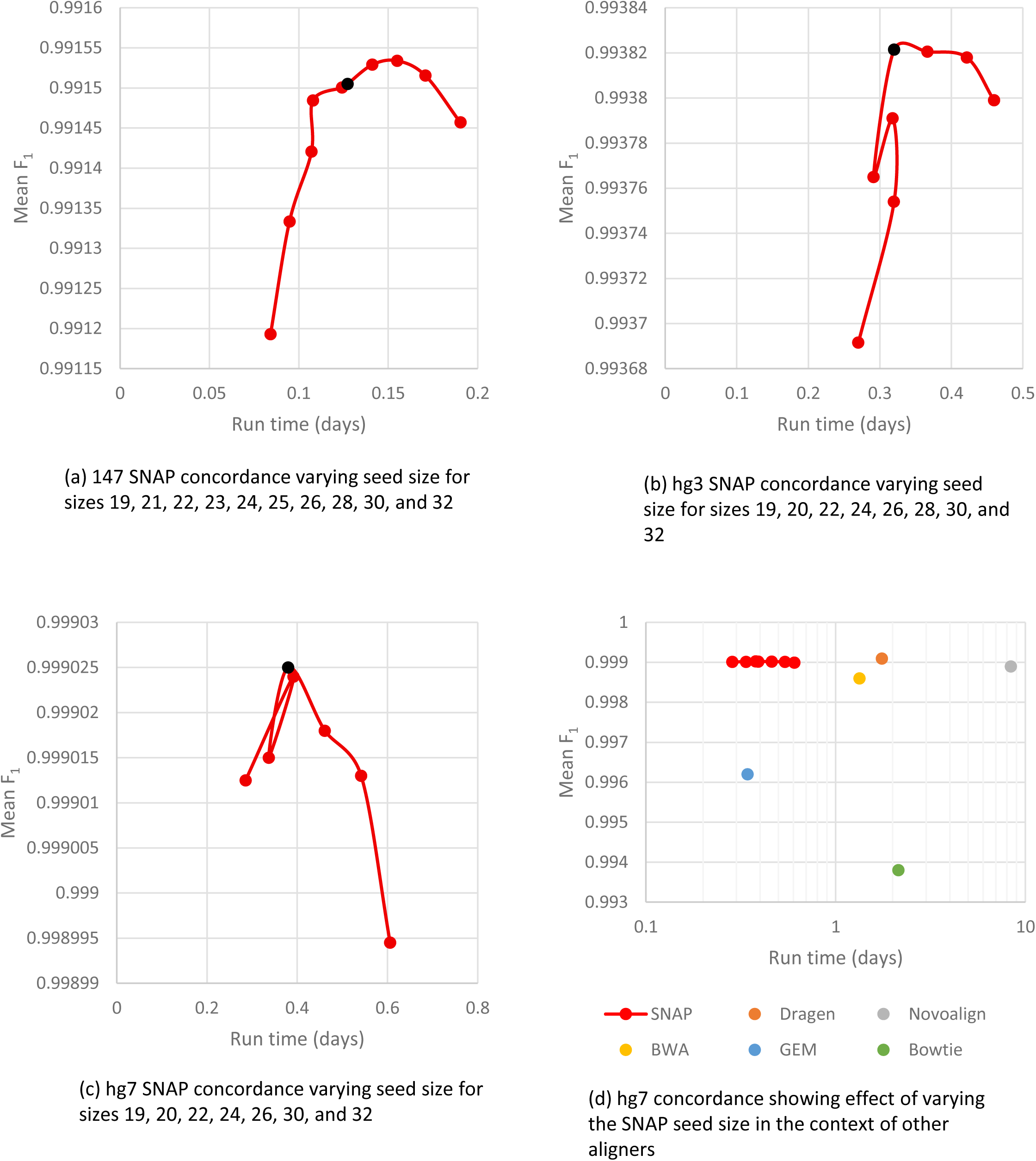
Effect of varying the seed size on SNAP concordance and performance. The black dots represent the default seed size of 24. In each case, the slowest point is seed size 19, and the line connects the runs in seed size order. It’s notable that while the performance effect can be about 2x between the extremes that the change in concordance is small, the largest being .00034 for 147 which is the sample with the smallest read length. This difference is dwarfed by the differences between aligners.

## Footnotes

1 https://1drv.ms/u/s!AhuEg_0yZD86hdE7OBoyZMWTiY2XJQ?e=JMyF2T

2 https://1drv.ms/u/s!AhuEg_0yZD86hdE8clsJJCAgXxSn7g?e=rS0P9d

3 While “FASTQ” is written in all capital letters, it is not an acronym but is simply the name of the file format.

